# Acclimation of *Dunaliella salina* to Olive Mill Wastewater: Oxidative Stress, Genotoxicity and the Role of Light Availability

**DOI:** 10.64898/2026.11.25.769106

**Authors:** Filippo Drigo, Lorenzo Mollo, Fabio Marcheggiani, Luca Tiano, Alessandra Norici

## Abstract

Freshwater resources are limited, and their availability is threatened by global climate change. Microalgae-based wastewater treatment enables water recovery for human activities, as microalgae can grow in wastewater while simultaneously removing contaminants within a circular economy framework. An example is OMWW. Thus, we tested the early and prolonged responses to tyrosol, *p*-coumaric acid, caffeic acid, their mixture and 10% OMWW on non-acclimated *Dunaliella salina* (Chlorophyta), analysing the effects through the DPPH and Comet assays. *D. salina* exhibited transient physiological damage after 1 day of exposure, whereas no detrimental effects were detected after 8 days, indicating the ability to withstand the toxicity of phenolic compounds and OMWW. Afterwards, we characterised the growth of acclimated *D. salina* cells to 10% OMWW at 100 and 300 μmol photons m⁻²s⁻¹, analysing the growth rate, pigment content, macromolecular composition and biomass reduction index. We observed that *D. salina* was able to grow similarly to the control in 10% OMWW when exposed to 300 μmol photons m⁻²s⁻¹, with a similar chl*a*/chl*b* ratio and carotenoids, a higher amount of proteins and carbohydrates and a more reduced biomass. In conclusion, *D. salina* demonstrated a high degree of tolerance to OMWW and high acclimatation capability. Moreover, this study indicates that growth of *D. salina* in OMWW is strongly light-dependent.

**Highlights:**

- Exposure to OMWW causes a transient oxidative and genotoxic stress response in *D. salina*.
- *D. salina* exhibits a remarkable acclimation capacity, mitigating the initial harmful effects of OMWW over time.
- Tyrosol is likely one of the main factors underlying the early toxicity observed following exposure.
- Light availability strongly influences the success of *D. salina* cultivation in OMWW.
- Although OMWW improves biomass biochemical composition, *D. salina* shows limited capacity for phenolic compound removal.

## 1. Introduction

Globally, only 1% of water resources are available for human consumption (e.g., drinking water, irrigation, power generation) (Bănăduc *et al.,* 2022), and this limited resource is also needed for many industrial activities, including pharmaceuticals, electronics, food, agrochemicals, and the oil and gas sector (Abdelfattah *et al.,* 2023). As a consequence, freshwater consumption and availability represent a major and growing issue for global stability (Abdel-Raouf *et al.,* 2012; Bănăduc *et al.,* 2022), further exacerbated by human-induced climate change, including increased temperatures and more frequent and intense heatwaves (Perkins *et al.,* 2012; Bănăduc *et al.,* 2022), as well as by water-intensive activities such as agriculture and data centers (Privette *et al.,* 2026). Strategies to recover and recycle water resources are therefore urgently needed.

One such strategy is wastewater treatment. Wastewater refers to all types of liquid discharge deriving from human activities (e.g., urban facilities, textile industries, livestock farms, agricultural processing plants). If not adequately treated, the high concentrations of organic and inorganic compounds it contains can compromise ecosystem health and impact human health (Akpor *et al.,* 2011; Liu and Yildiz, 2018; Ilyas *et al.,* 2019; Abdelfattah *et al.,* 2023). Among the different types of industrial wastewater, Olive Mill Wastewaters (OMWW) represent a serious environmental pollutant in Mediterranean countries with a high production of olive oil (Paraskeva and Diamadopoulos, 2006; Markou *et al.,* 2012). Their environmental footprint was demonstrated by Pavlidou and co-authors (2014), who showed that the ecological status of rivers and coastal areas does not return to optimal levels before 5 months from their release. This impact is largely attributable to their low pH, high concentration of suspended solids, high chemical and biological oxygen demand (COD and BOD), and high concentrations of phenolic compounds (Paraskeva and Diamadopoulos, 2006; Danellakis *et al.,* 2011).

Phenolic compounds, or polyphenols, are a large and structurally diverse group of plant secondary metabolites that are naturally abundant in olive-derived matrices and are therefore major constituents of OMWW (Abbas *et al.,* 2017). These molecules display a dual biological nature: at low concentrations they are widely recognized for their antioxidant and health-promoting properties (Abbas *et al.,* 2017), while at higher concentrations, or in different biological systems, they can behave as pro-oxidants, generating reactive oxygen species and causing oxidative and genotoxic damage (Scalbert *et al.,* 2005). This duality is central to understanding both the toxicity of OMWW towards aquatic organisms and the potential biological value of the phenolic-rich biomass that can be produced when these compounds are metabolized rather than simply degraded.

In particular, phenolic compounds are known to affect important primary producers such as microalgae, a diverse group of unicellular photosynthetic organisms. As mentioned above, they can have a pro-oxidant nature (Scalbert *et al.,* 2005; Abbas *et al.,* 2017), but they can also be metabolized by microalgae as a carbon source (Lindner and Pleissner, 2019). The effects of these compounds depend on their chemical nature, their concentration, and the species considered (Awad *et al.,* 2020; Mollo *et al.,* 2023): they can inhibit or promote microalgal growth (Lindner and Pleissner, 2019; Awad *et al.,* 2020), and can promote the synthesis of specific compounds at specific concentrations (Zou *et al.,* 2023; Sun *et al.,* 2024).

In fact, microalgae can be used in wastewater treatment applications, using these waters as a growth medium (Abdel-Raouf *et al.,* 2012), as demonstrated by previous applications (Liu and Yildiz, 2018; Abdelfattah *et al.,* 2023), within the frame of a circular economy. This strategy offers several potential advantages: (1) it reduces the environmental impact of wastewater; (2) it reduces the amount of freshwater needed for microalgae cultivation; (3) it reduces the costs associated with both wastewater treatment and microalgae cultivation (Abdel-Raouf *et al.,* 2012; Surkatti & Al-Zuhair, 2018; Calijuri *et al.,* 2022). Thanks to their metabolic adaptability (Lindner and Pleissner, 2022), microalgae can remove heavy metals and some toxic organic compounds and can reduce the nitrogen and phosphorus loads of wastewaters (Abdel-Raouf *et al.,* 2012; Liu and Yildiz, 2018). Additionally, the resulting biomass is an important source of bioactive compounds (Santhosh *et al.,* 2016; Liu and Yildiz, 2018), relevant for feedstock, biofuel, and pharmaceutical applications (Santhosh *et al.,* 2016; Cicci *et al.,* 2024).

Among the available microalgal strains, *Dunaliella salina* (Chlorophyta) is of particular interest (Feng *et al.,* 2020). One of its most relevant features is its halophilic nature, which further reduces the need for freshwater resources during cultivation (Liu and Yildiz, 2018; De Souza Celente *et al.,* 2024). Beyond its well-known ability to accumulate β-carotene (Liu and Yildiz, 2018), *D. salina* is increasingly recognized as a versatile biotechnological platform: depending on the strain and culture conditions, it can also accumulate other carotenoids, glycerol, lipids, and proteins, supporting applications in the food, feed, cosmetic, and pharmaceutical sectors (Barbosa *et al.,* 2023; Prajapat *et al.,* 2026). This species has also been successfully used in wastewater treatment applications (Hu *et al*., 2024; Barreto *et al.,* 2025), obtaining interesting results in terms of biomass characteristics. Lastly, a previous screening of microalgal tolerance to the most abundant phenolic compounds in OMWW (Mollo *et al.,* 2023) demonstrated *D. salina* resistance to these molecules.

Within this framework, we assessed the initial effects of three phenolic compounds (tyrosol, *p*-coumaric acid, and caffeic acid), tested individually and in combination, together with OMWW, on non-acclimated *D. salina* cells using DPPH and Comet assays. Additionally, because *D. salina* exhibited only a transient oxidative and genotoxic stress response, we examined whether acclimated cells grown in 10% diluted OMWW could benefit from a threefold increase in irradiance (300 μmol photons m⁻² s⁻¹), relative to the control light intensity of 100 μmol photons m⁻² s⁻¹.

## 2. Materials and methods

Due to the high content of suspended solids, organic matter, and the presence of indigenous microbial and fungal populations, Olive Mill Wastewater (OMWW) was pre-treated via sequential filtration. The effluent was filtered through nitrocellulose membranes down to a 0.2 µm pore size to remove particulate matter and reduce the microbial load, rendering the OMWW suitable for microalgal growth. Filtered OMWW was stored at −20 °C until use.

### 2.1 Toxicity Assessment of Phenolic Compounds and OMWW

The toxic effects of specific phenolic compounds (tyrosol, *p*-coumaric acid, and caffeic acid, tested both individually and as a mixture) and 10% (v/v) OMWW were evaluated on non-acclimated cultures of *Dunaliella salina* (isolated from the Margherita di Savoia saltpans, Italy) (Mollo *et al*., 2023). Experimental cultures were inoculated in triplicate in Erlenmeyer flasks with a 100 mL working volume at an initial density of 10^5^ cells mL^−1^, using parent cultures maintained in AMCONA medium (Fanesi *et al.,* 2014) and harvested during the exponential growth phase.

Individual phenolic treatments and their mixture were performed at 100 μmol photons m^−2^ s^−1^ (condition called Low Light, LL) at 20 ± 2 °C. Phenolic concentrations were set at one-fifth (1/5) of the levels commonly found in OMWWs and reported by Deeb *et al*. (2014), corresponding to 42.12 mg L^−1^ for tyrosol, 23.42 mg L^−1^ for *p*-coumaric acid, and 28.08 mg L^−1^ for caffeic acid. The mixture treatment contained a total of 93.62 mg L^−1^ of phenols, corresponding to the sum of the individual compounds.

The 10% OMWW treatment was conducted under a High Light regime (HL, 300 μmol photons m^−2^ s^−1^ at room temperature) due to the high turbidity of the growth medium (Supplementary Fig. 1).

All experimental sets were maintained under a 24-h continuous photoperiod and compared against their respective LL and HL environmental control groups (cultures grown in AMCONA medium). Algal growth and viability were monitored daily using a Casy TT automatic cell counter (Innovatis AG, Reutlingen, Germany) for 8 days, corresponding to the end of the exponential phase as observed in Mollo *et al.,* (2023). As in a continuous industrial process and in similar experiments, all the analysis were carried out only during the exponential phase (Liu and Yildiz, 2017; Awad *et al.,* 2020).

#### 2.1.1 Antioxidant capacity (DPPH Assay)

To evaluate the oxidative stress response, antioxidant capacity was measured 2- and 7-days post-inoculum using the 2,2-diphenyl-1-picrylhydrazyl (DPPH•) radical scavenging assay. For each sample, a biomass volume corresponding to a final concentration of 0.5 mg mL^−1^ in ethanol was collected by centrifugation (3260xg, 15 min, 4 °C). Cells were mechanically disrupted using a pestle to release intracellular antioxidants and stored in darkness at −20 °C. The assay was performed by reacting 100 µL of ethanolic extract with 900 µL of a 0.06 mM DPPH• methanolic solution. After 15 minutes of incubation, absorbance was recorded at 515 nm (UV-1900i, Shimadzu Corp.). The Percentage of Inhibition (PI) of DPPH• was calculated according to Assunção *et al*. (2017):

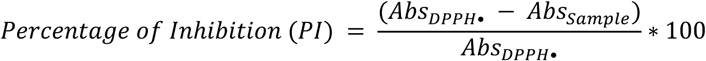

Where Abs_DPPH•_ represents the control (900 µL DPPH• + 100 µL ethanol) and Abs_blank_ accounts for the specific absorbance of the extract (100 µL extract + 900 µL methanol) to eliminate interference from high carotenoid content.

#### 2.1.2 Genotoxic effects assessment (Comet assay)

Genotoxic effects were evaluated using the Alkaline Comet Assay (Single Cell Gel Electrophoresis) 1- and 8-days post-inoculum. For each treatment, approximately 8×10^3^ cells were harvested by centrifugation (600xg, 15 min, 4 °C). Positive controls for DNA damage were prepared by exposing cells for 15 minutes to two hydrogen peroxide solutions (H_2_O_2_: 0.75, 1.5 mM) prepared in growth medium to avoid osmotic shock.

Both experimental samples and positive controls were washed with 0.5 M PBS and resuspended in 100 µL of 1% Low Melting Point Agarose (LMA) at 37 °C. Aliquots of 35 µL (approx. 2.8×10^3^ cells) were then solidified on pre-chilled slides. Cell lysis was performed at 4°C for 1 hour in a specific lysis buffer, followed by DNA denaturation in an alkaline electrophoretic buffer for 20 minutes. Electrophoresis was carried out at 20 V for 20 minutes. Subsequently, slides were neutralized, dehydrated in 70% methanol, and dried at 60 °C. DNA was stained using SYBR™ Gold Nucleic Acid Gel Stain (S11494, Thermofisher U.S.A) at 1x diluited in PBS, from stock solution 10,000X concentration in DMSO,, and images were acquired via Lionheart FX Automated Microscope (Biotek, U.S.A.) (12 images per well). DNA damage was quantified using image analysis software to determine four primary indices: Tail Length (TL), Tail Intensity (TI) and the Tail Moment (TM) which combines TL and TI (De Bernardi *et al*., 2022; Marini *et al.,* 2024).

### 2.2 Algal Acclimation, Cultivation and Biomass Analysis in OMWW

To characterize *D. salina* growth in 10% OMWW (v/v), we performed cultivation trials of acclimated cells for at least 5 generations in Erlenmeyer flasks with a 100 mL working volume. Two distinct environmental regimes were evaluated under a 24-hour continuous photoperiod: (1) Low Light (LL) - 100 μmol photons m^−2^ s^−1^ at 20 ± 2 °C; (2) High Light (HL) - 300 μmol photons m^−2^ s^−1^ at RT (Room Temperature). For each regime, cultures were grown in AMCONA medium supplemented with 10% (v/v) filtered OMWW. Control groups were maintained in pure AMCONA medium under the same LL and HL conditions to provide a baseline for physiological comparisons. Cultures were inoculated at an initial concentration of 10^5^ cells mL^−1^ using parent cultures previously acclimated to the saline-wastewater mixture and maintained in the exponential growth phase.

Algal growth was monitored daily by counting cells using an automatic cell counter (Casy TT, Innovatis AG, Reutlingen, Germany). Monitoring continued until the end of the exponential phase, which occurred after 7 days for both light conditions.

#### 2.2.1 Quantification of phenolic compounds

Total phenolic content was quantified spectrophotometrically every two days throughout the experiment using a modified Folin-Ciocalteu method, as described in Mollo *et al*. (2024).

Briefly, 125 µL of culture was centrifuged, and the resulting supernatant was reacted with 125 µL of Folin-Ciocalteu reagent under vigorous stirring. After 6 minutes, 2.25 mL of 4% NaHCO_3_ was added, and the mixture was incubated in the dark for 90 minutes. Absorbance was measured at 760 nm (UV-1900i, Shimadzu Corp.), and concentrations were determined using a gallic acid standard calibration curve (0-100 mg L^−1^). Results were expressed as a percentage of the initial phenolic concentration.

#### 2.2.2 Pigment Extraction and Quantification

Photosynthetic pigments were analyzed during the exponential phase. For each biological replicate, two 1.5 mL culture aliquots were collected as instrumental replicates. Samples were centrifuged at 13,000 rpm for 10 minutes, and the resulting pellet was washed with 0.5 M ammonium formate (NH_4_HCO_2_) to remove medium interference. After a second centrifugation, pigments were extracted by resuspending the pellet in 90% acetone (CH_3_COCH_3_) and storing the samples at −20 °C in darkness until the pellet appeared colourless. Absorbance spectra (350–750 nm, 5 nm steps) were recorded using a UV-1900i spectrophotometer (Shimadzu Corp.) with quartz cuvettes. Chlorophyll *a* (chl *a*), chlorophyll *b* (chl *b*), and total carotenoids were calculated according to equations proposed by Singh *et al*. (2019). Pigment content was expressed as pg cell^−1^.

#### 2.2.3 Protein extraction and quantification

Cellular protein content was determined during the exponential phase, following the Lowry method (Lowry *et al.,* 1951) as modified by Peterson (1977). For each biological replicate, two 1.5 mL aliquots were centrifuged (13,000 rpm, 10 min) and the pellets washed with 0.5 M ammonium formate. Following, the pellet was resuspended in 500 µL of 1% (w/v) SDS and 0.1 M NaOH and vigorously vortexed. Subsequently, 500 µL of Reagent A was added; after 10 minutes, 250 µL of Reagent B (Folin-Ciocalteu) was introduced, and the mixture was incubated in the dark for 30 minutes. Absorbance was measured at 750 nm (UV-1900i, Shimadzu Corp.). Quantification was performed using a bovine serum albumin (BSA) calibration curve (0-100 μg). Final values were normalized to cell density and expressed as pg cell^−1^.

#### 2.2.4 FTIR Analysis and Macromolecular Semiquantification

Macromolecular composition was analyzed during the exponential phase, using a Tensor 27 FTIR spectrometer (Bruker Optics, Ettlingen, Germany), with sample preparation and spectra acquisition following the protocols of Domenighini and Giordano (2009).

Bands were assigned to cellular components (protein, carbohydrates and lipids) following Giordano *et al*. (2001). After spectral deconvolution, band integrals were used to estimate the relative fractions of carbohydrates, lipids and proteins, using OPUS 6.5 (Bruker Optik GmbH). Semi-quantification of cellular pools from FTIR spectra followed the procedure described in Palmucci *et al*. (2011). To facilitate comparison between different growth regimes, results expressed in arbitrary units (AU) were further normalized by setting the values of the control condition to 1.

#### 2.2.5 Biomass reduction index

The biomass reduction index was determined via FTIR spectroscopy to evaluate changes in the relative proportions of CH bond in CH*_n_* groups. Samples were collected during the exponential phase and prepared following the same procedures described for macromolecular characterization (see 2.1.4). For this analysis, the spectral deconvolution was focused on the areas of specific CH groups to evaluate biomass reduction: CH (2920 cm^−1^), CH_2_ (2950 cm^−1^), CH_3_ (3015 cm^−1^) groups.

The reduction index was calculated according to Palmucci *et al*. (2011) using the following ratio:

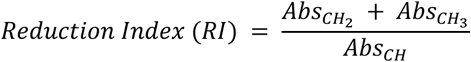

It is important to note that this value does not represent an absolute reduction level but indicates a shift in the relative abundance of these groups. Results were normalized by setting the values of the control condition to 1.

### 2.3 Statistical analysis

All data are expressed as mean ± standard deviation (SD) of three independent biological replicates. For FTIR, Comet assay, and spectrophotometric measurements, two instrumental replicates were performed for each biological sample. Statistical significance was evaluated using R-studio software (v 4.5.1). Depending on the experimental design, differences between means were determined using Student’s t-test, one-way ANOVA or two-way ANOVA, both followed by Tukey’s post-hoc test for multiple comparisons. The significance threshold was set at 5% (*p*-value < 0.05). Detailed ANOVA summary tables are provided in Supplementary Tables 1, 2, 3.

## 3. Results

### 3.1 Oxidative effects of phenolic compounds and OMWW on short-term exposed *D. salina* cells

#### 3.1.1 Growth response

Figure 1 shows the growth response of *D. salina* cultures exposed to different phenolic compounds. Cell growth rates under all treatment conditions were comparable to those of the respective control culture as shown in Table 1. The only significant difference was observed in the cell concentration after 8 days of exposure to the mixture of phenols (Table 1).

**Figure 1.**
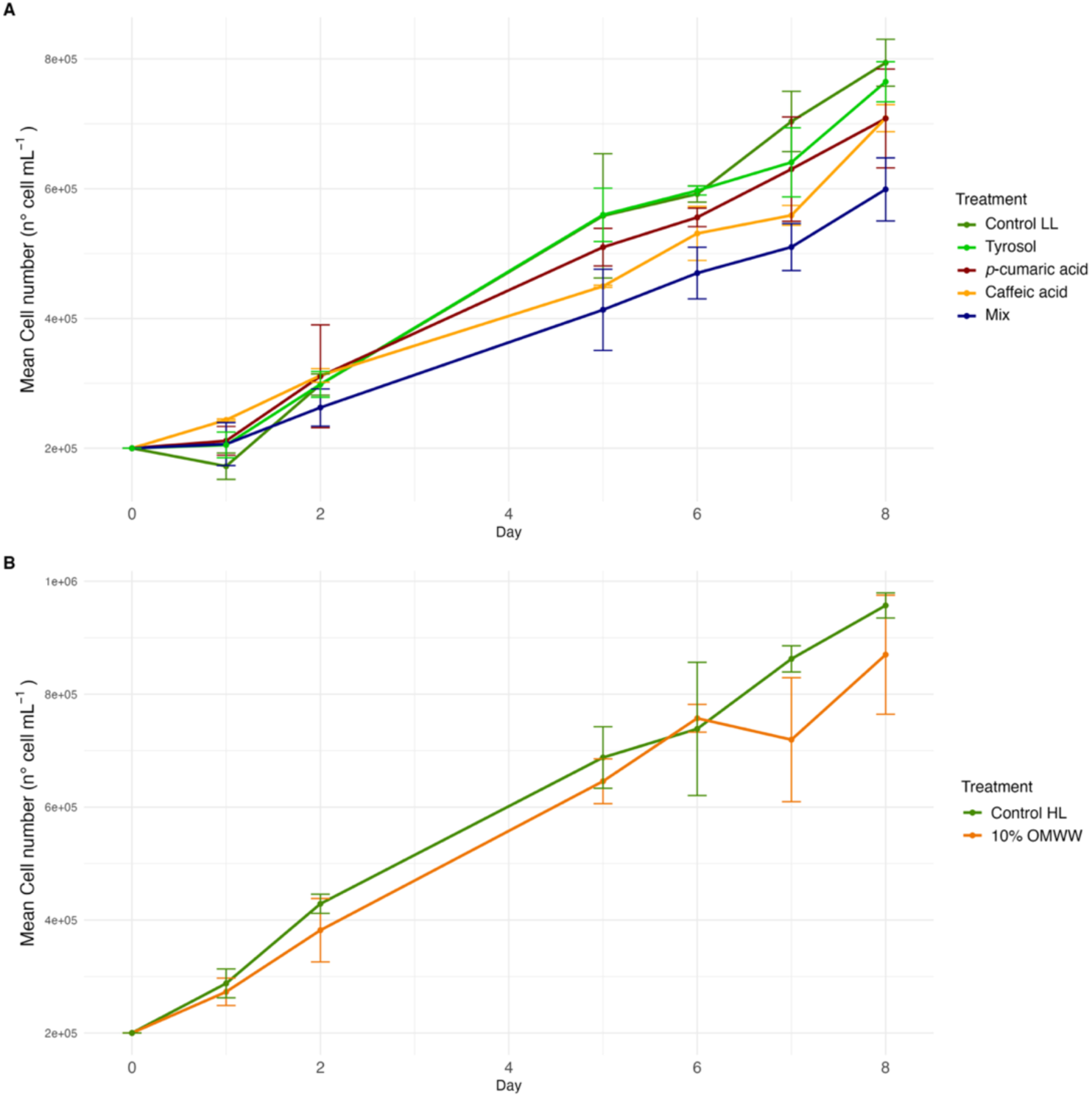
Growth curve of *D. salina* exposed to phenolic compounds and 10% OMWW. Concentration of *D. salina* in control condition at LL (green), exposed to tyrosol (light green), *p*-coumaric acid (red), Caffeic acid (yellow), Mix (blue), control condition at HL (orange) and in 10% OMWW (red) expressed as Mean cell number (n° cell mL^−1^) ± SD per day.

**Table 1.**
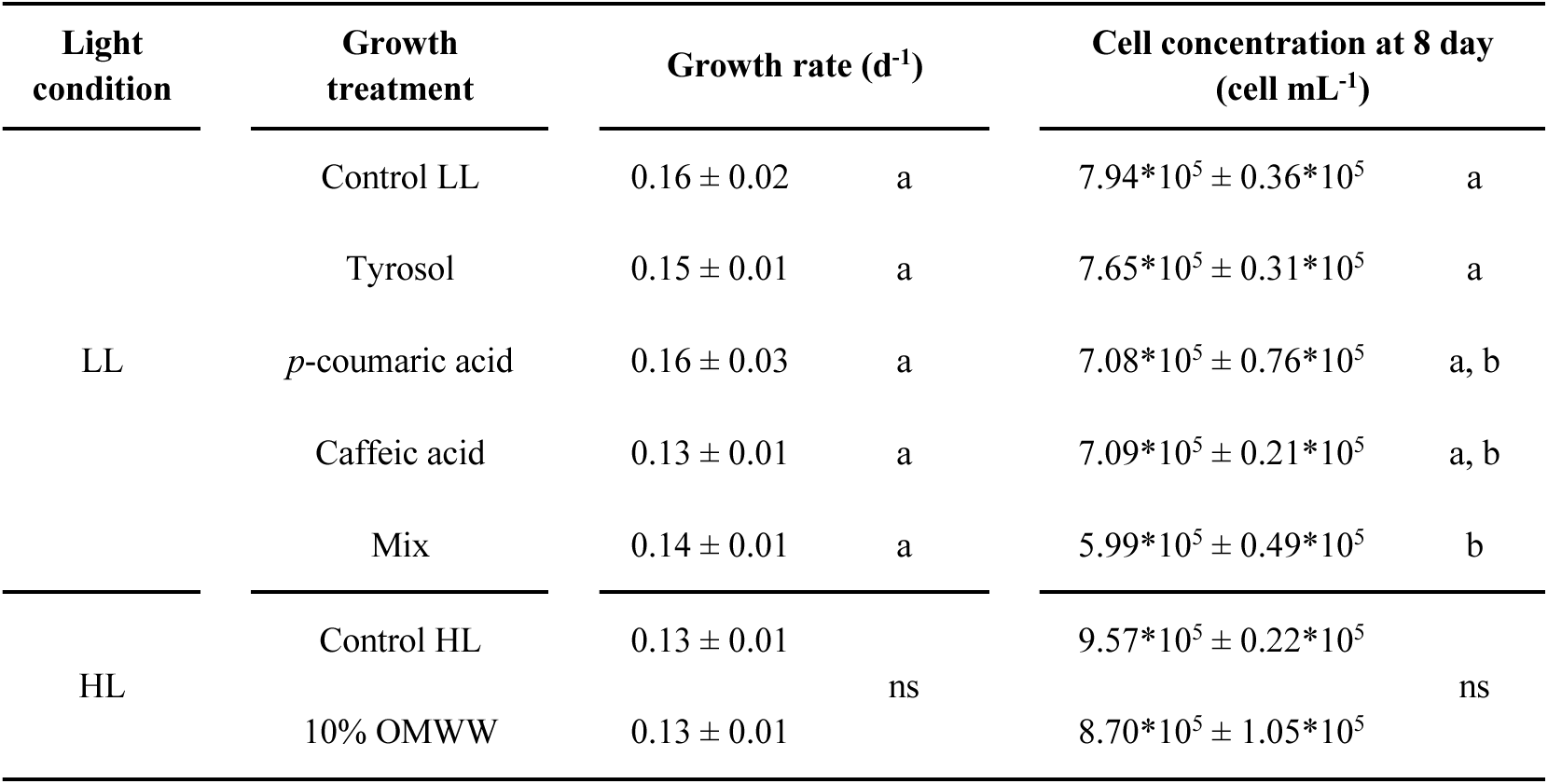
Growth rate and Cell concentration after 8 days of exposure of *D. salina* growth under different treatments. Data are reported as mean ± standard deviation. One-way ANOVA was used to evaluate differences among Control LL, tyrosol, *p*-coumaric acid, Caffeic acid and Mix conditions. Values followed by different letters are significantly different (adjusted p-value < 0.05), additional details available in Supplementary Table 1. Student’s t-test was performed to evaluate differences between Control HL and 10% OMWW conditions.

| **Light condition** | **Growth treatment** | **Growth rate (d<sup>-1</sup>)** |  | **Cell concentration at 8 day (cell mL<sup>-1</sup>)** |  |
|---|---|---|---|---|---|
| LL | Control LL | 0.16 ± 0.02 | a | 7.94*10<sup>5</sup> ± 0.36*10<sup>5</sup> | a |
|  | Tyrosol | 0.15 ± 0.01 | a | 7.65*10<sup>5</sup> ± 0.31*10<sup>5</sup> | a |
|  | *p*-coumaric acid | 0.16 ± 0.03 | a | 7.08*10<sup>5</sup> ± 0.76*10<sup>5</sup> | a, b |
|  | Caffeic acid | 0.13 ± 0.01 | a | 7.09*10<sup>5</sup> ± 0.21*10<sup>5</sup> | a, b |
|  | Mix | 0.14 ± 0.01 | a | 5.99*10<sup>5</sup> ± 0.49*10<sup>5</sup> | b |
| HL | Control HL | 0.13 ± 0.01 | ns | 9.57*10<sup>5</sup> ± 0.22*10<sup>5</sup> | ns |
|  | 10% OMWW | 0.13 ± 0.01 |  | 8.70*10<sup>5</sup> ± 1.05*10<sup>5</sup> |  |

#### 3.1.2 Antioxidant activity

As shown in Fig. 2A, after 48h hours of exposure, antioxidant activities were identical among treatments and control except for the 10% OMWW treatment showing a higher inhibition percentage. After 7 days of exposure, cultures exposed to caffeic acid and the mixture of phenolic compounds had the lowest DPPH inhibition activity while all other date were similar. Considering the variation in inhibition percentage from the 2^nd^ day to the 7^th^ day (Fig. 2B), only in the tyrosol growth condition a higher inhibition percentage was observed while a significantly lower percentage from the initial time point was observed in the treatments containing caffeic acid, the phenolic mixture and 10% OMWW.

**Figure 2.**
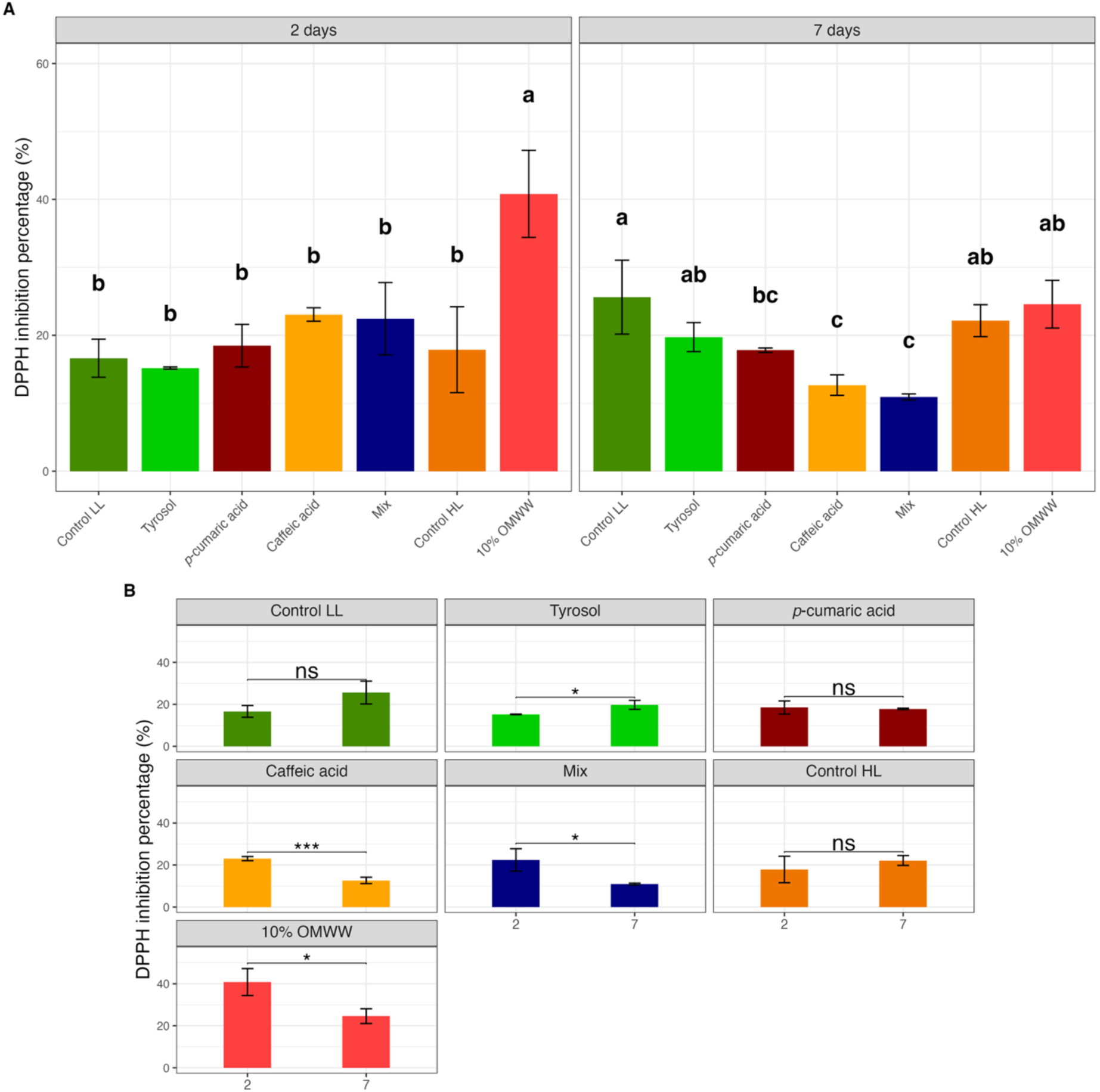
Inhibition percentage of DPPH• in *D. salina* exposed to phenolic compounds and 10% OMWW at different time points. DPPH• inhibition percentage in *D. salina* in control conditions at LL (green), exposed to tyrosol (light green), *p*-coumaric acid (bordeaux), Caffeic acid (yellow), Mix (blue), control conditions at HL (orange) and in 10% OMWW (red). **(A)** Comparison among the different growth conditions after 2 (left) and 7 (right) days post-inoculum. Significant differences are represented by different letters, more details available in Supplementary Table 1. **(B)** Comparison between the Inhibition percentage after 2- and 7-days post-inoculum for the different growth conditions. Asterisks indicate significant differences: \**p*-value<0.05, \*\*\**p*-value<0.001.

#### 3.1.3 Genotoxic effects

After 24 hours of exposure the one-way ANOVA analysis of Tail Length (TL) did not detect significant differences between the different growth treatments (Fig. 3A), the same occurred after 8 days of exposure. Conversely, the comparison between the two time-points indicated a significant reduction of TL only in the treatments containing tyrosol (*p-value*: 3.81*10^−5^), *p*-coumaric acid (*p-value*: 0.043), and 10% OMWW (*p-value*: 0.0003) (Fig. 3B).

**Figure 3.**
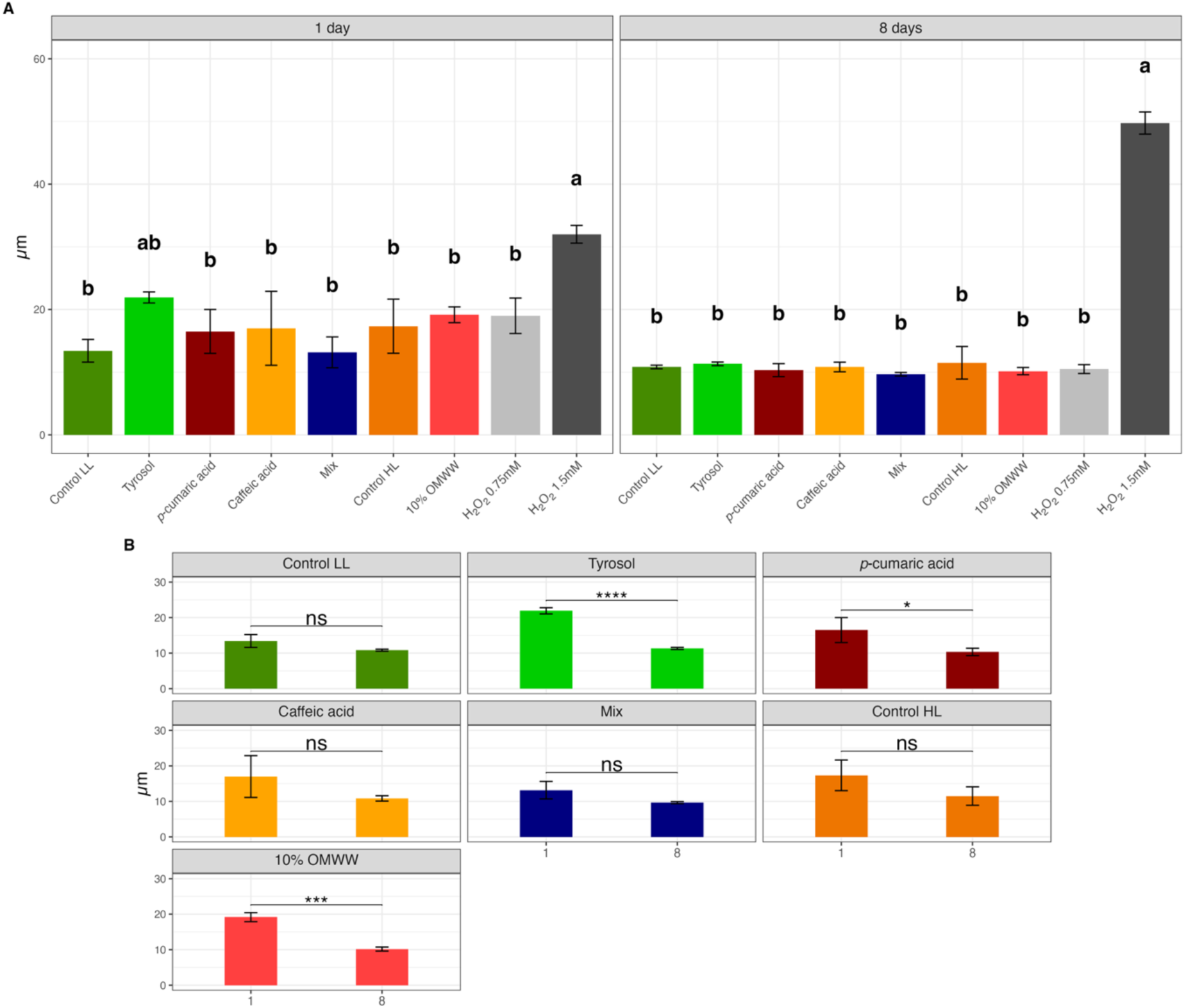
Tail length in *D. salina* exposed to phenolic compounds and 10% OMWW at different time points. Tail length (µm) in *D. salina* in control conditions at LL (green), exposed to tyrosol (light green), *p*-coumaric acid (bordeaux), Caffeic acid (yellow), Mix (blue), control conditions at HL (orange), 10% OMWW (red), H_2_O_2_ 0.75 mM (gray) and H_2_O_2_ 1.5 mM (dark gray). **(A)** Comparison among the different treatments after 1 (left) and 8 (right) days post-inoculum. Significant differences are represented by different letters, more details available in Supplementary Table 1. **(B)** Comparison between the TL after 1- and 8-days post-inoculum for the different treatments. Asterisks indicate significant differences: \**p*-value<0.05, \*\*\**p*-value<0.001, \*\*\*\**p*-value<0.0001.

Differently from TL, after 24 hours of exposure, the Tail Intensity (TI, which measures the % of DNA in the tail) of the 10% OMWW exposure was significantly higher than that of Control LL, *p*-coumaric acid and Mix conditions (Fig. 4A). No differences were observed after 8 days, with the exception of the 10% OMWW culture, whose TI was still significantly higher than that of CTR LL and Mix treated cultures. The comparison between the two time-points indicated a significant decrease in the Control LL, tyrosol and 10% OMWW treatments. In particular, the last two decreased by ∼15% from the first to the second time point (Fig. 4B).

**Figure 4.**
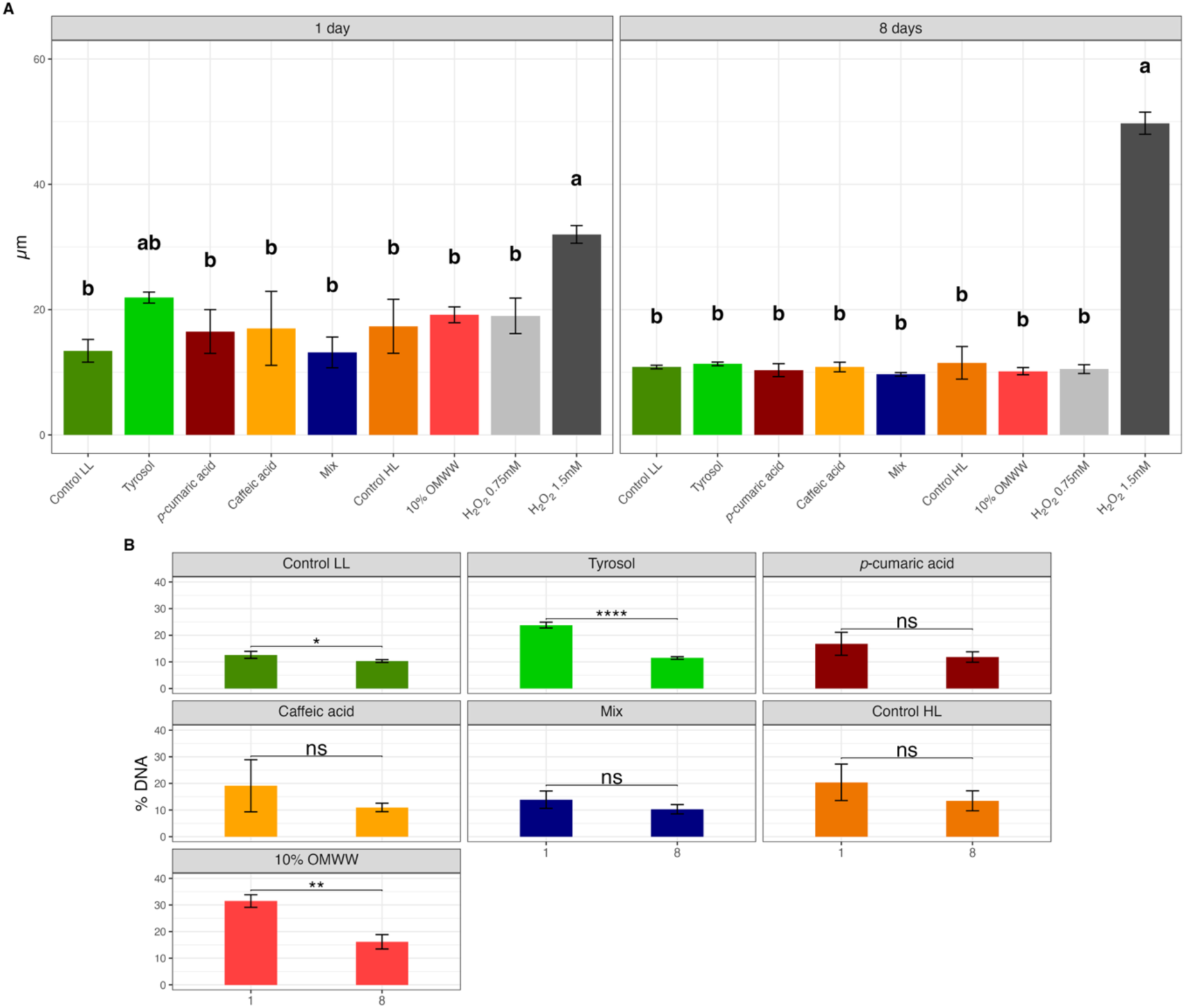
Tail intensity in *D. salina* exposed to phenolic compounds and 10% OMWW at different time points. Tail intensity (% DNA) of *D. salina* in control cultures at LL (green), exposed to tyrosol (light green), *p*-coumaric acid (bordeaux), Caffeic acid (yellow), Mix (blue), control cultures at HL (orange), 10% OMWW (red), H_2_O_2_ 0.75 mM (gray) and H_2_O_2_ 1.5 mM (dark gray). **(A)** Comparison among the different growth treatments after 1 (left) and 8 (right) days post-inoculum. Significant differences are represented by different letters, more details available in Supplementary Table 1. **(B)** Comparison between the TI after 1- and 8-days post-inoculum for the different growth conditions. Asterisks indicate significant differences: \**p*-value<0.05, \*\**p*-value<0.01, \*\*\*\**p*-value<0.0001.

Lastly, the one-way ANOVA analysis of Tail Moment (TM) indicated a significantly higher value in the 10% OMWW culture compared to the value of Control LL, *p*-coumaric acid, Caffeic acid, Mix and Control HL cultures after 24h of exposure. After 8 days no significant differences were detected (Fig. 5A). Considering the differences between the two time points for each treatment, we measured a significant decline of TM in the Control LL culture and in the cultures exposed to tyrosol and 10% OMWW (Fig. 5B).

**Figure 5.**
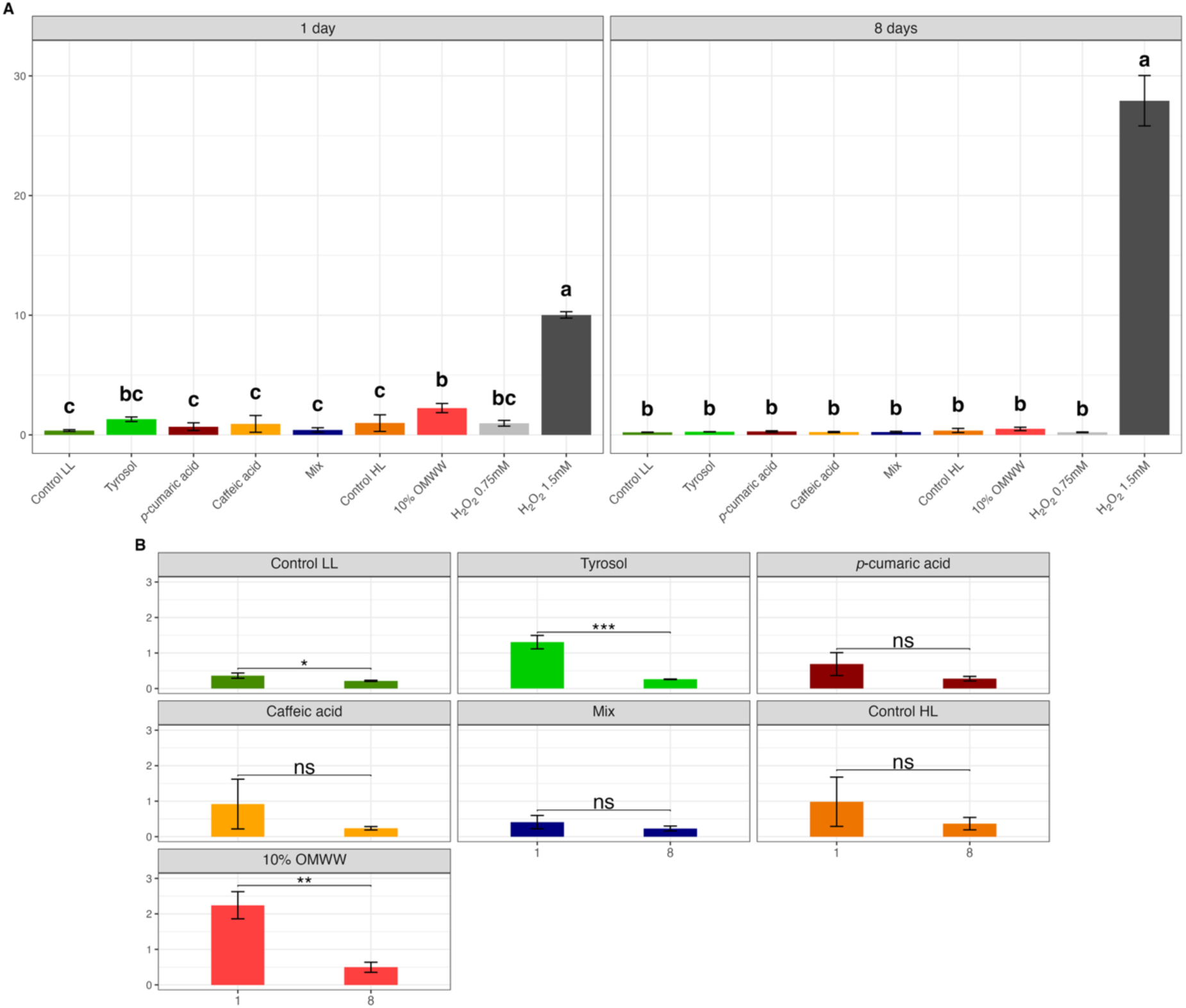
Tail moment in *D. salina* exposed to phenolic compounds and 10% OMWW at different time points. Tail moment of *D. salina* in control cultures at LL (green), exposed to tyrosol (light green), *p*-coumaric acid (bordeaux), Caffeic acid (yellow), Mix (blue), control cultures at HL (orange), 10% OMWW (red), H_2_O_2_ 0.75 mM (gray) and H_2_O_2_ 1.5 mM (dark gray). **(A)** Comparison among the different growth treatments after 1 (left) and 8 (right) days post-inoculum. Significant differences are represented by different letters, more details available in Supplementary Table 1. **(B)** Comparison between the TM after 1- and 8-days post-inoculum for the different growth conditions. Asterisks indicate significant differences: \**p*-value<0.05, \*\**p*-value<0.01, \*\*\**p*-value<0.001.

Overall, the two treatments with the highest genotoxic effects among the one tested resulted to be the tyrosol and 10% OMWW, especially in terms of TI and TM.

### 3.2 *D. salina* acclimated to OMWW based growth medium

#### 3.2.1 Growth

As shown by Fig. 6, cells of *D. salina* acclimated to 10% OMWW LL showed significantly lower growth rate, division time and maximum density than in control cells after 7 days of growth (Fig. 6A. 6C). Acclimation of the algae to higher light (HL) intensity resulted in faster growth, as shown in Figure 6B. Under this condition, the growth rate and division time (Figure 6D) were comparable to those of the control cultures, with significant differences observed only in the maximum cell density achieved. In both light regimes, treated cells had a smaller cell volume relative to control cells (LL: 1649 ± 66 fL in the control vs. 957 ± 44 fL in 10% OMWW; HL: 1661 ± 19 fL in the control vs. 1033 ± 2 fL in 10% OMWW). Despite the recovery in terms of growth parameters of the 10% OMWW HL grown culture, we did not observe any significant reduction of the phenolic concentration in the growth medium, as shown in Supplementary Figure 2.

**Figure 6.**
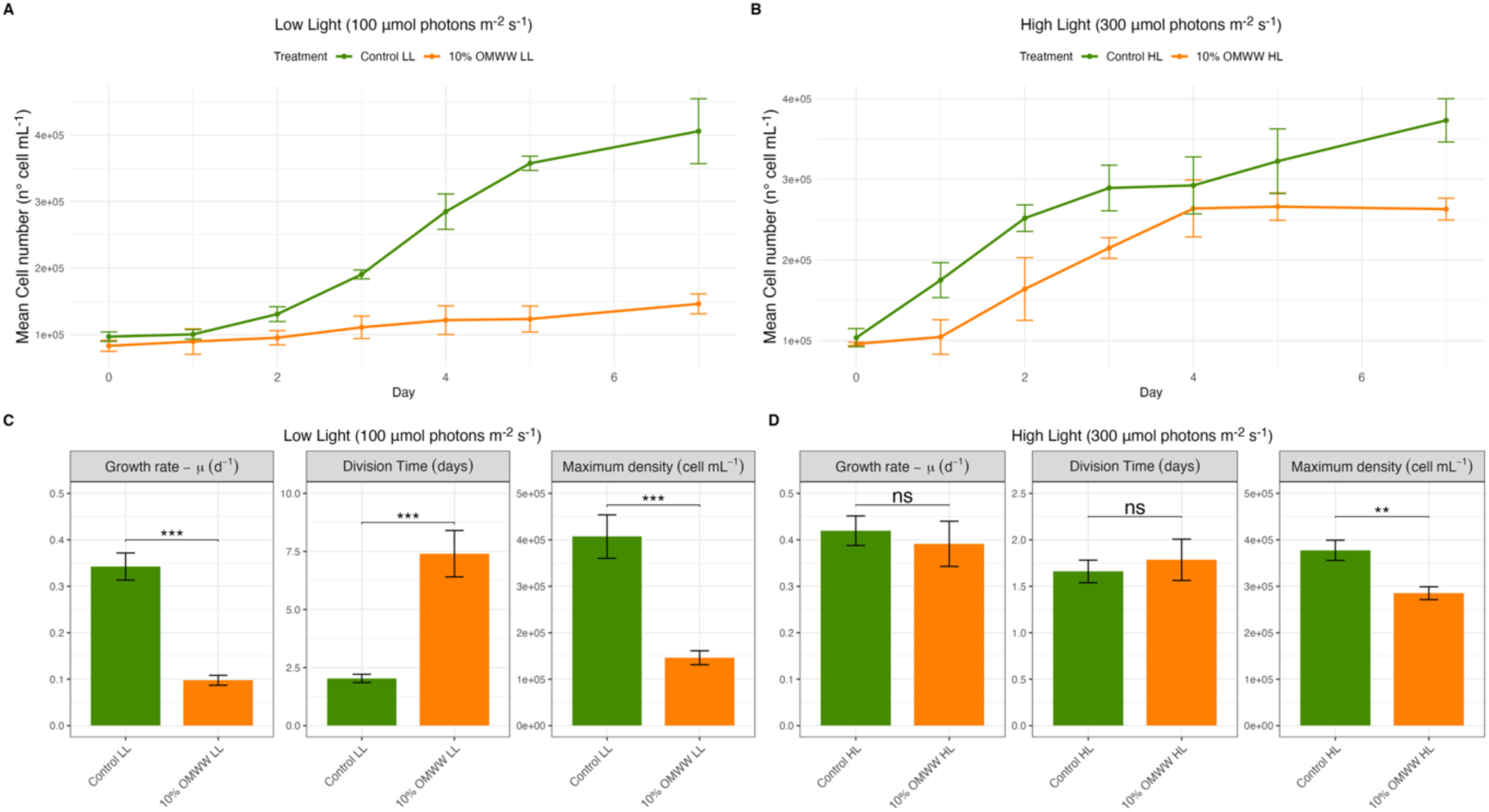
Growth characteristics of *D. salina* at Low Light (LL) and High Light (HL) in 10 % OMWW. (**A**) Growth curves of *D. salina* in control medium (green) and in 10% OMWW LL (orange). (**B**) Growth curves of *D. salina* in control medium (green) and in 10% OMWW HL (orange). (**C**) Growth rate (µ - d^−1^), Division Time (days) and Maximum Density (n° cell/mL) of *D. salina* in control (green) and 10% OMWW LL (orange). (**D**) Growth rate (µ - d^−1^), Division Time (days) and Maximum Density (n° cell/mL) of *D. salina* in control (green) and 10% OMWW HL (orange). Asterisks indicate significant differences: \**p*-value<0.05, \*\**p*-value<0.01, \*\*\**p*-value<0.001.

#### 3.2.2 Pigment content

Under LL, we measured a significantly lower content of chlorophylls (*a* and *b*) and total carotenoids in the treated cultures as compared to the contents in control cultures (Fig. 7A, B, D). Also the chl *a*/chl *b* ratio was significantly lower in the treated cultures (Fig. 7C).

**Figure 7.**
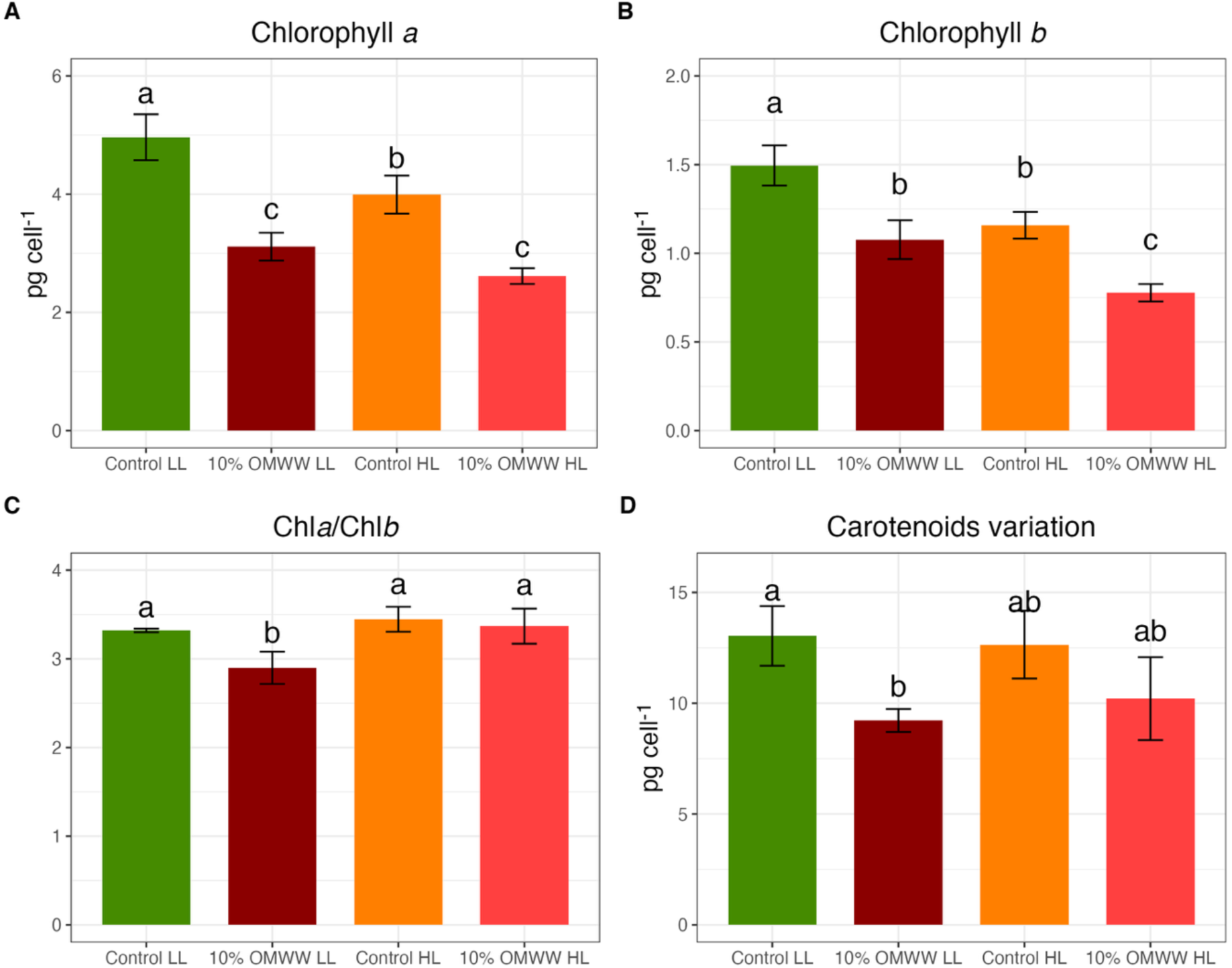
Pigments profiles of *D. salina* at Low Light (LL) and High Light (HL) in 10% OMWW. Quantification of (**A**) chlorophyll *a* (pg cell^−1^), (**B**) chlorophyll *b* (pg cell^−1^), (**C**) chlorophyll *a*/chlorophyll *b* and (**C**) total carotenoids (pg cell^−1^) of *D. salina* in LL control culture (green), 10% OMWW LL cultures (bordeaux), HL control cultures (orange) and 10% OMWW HL cultures (orange) during the exponential growth phase. Different letters represent statistical differences between groups resulting from two-way ANOVA analysis (factors: growth treatment and light regime).

Conversely, despite significant differences in the content of chl *a* and chl *b* between the control and 10% OMWW HL treatment (Fig. 7 A, B), the chl *a*/chl *b* ratio was not significantly different to the control, as for total carotenoids (Fig. 7C, D). All pigments were significantly influenced by the presence of 10% OMWW in the medium and the light regime, but not by their interaction, with only the exception of carotenoids where the growth in 10% OMWW was the only significant factor (Supplementary Table 2 and 3).

#### 3.2.3 Macromolecular profile

Under both light conditions, *D. salina* protein content per cell was significantly higher in the 10% OMWW grown culture compared to the control culture (Fig. 8A), with the difference being more pronounced at LL. At the same time, in the exponential phases of both light conditions, *D. salina* in 10% OMWW based medium showed a significantly more abundant carbohydrates (∼2 times at LL and ∼2.3 at HL) as compared to the control values (Fig. 8B). Regarding the lipid pool, we observed a significantly higher amount of lipids in the 10% OMWW LL (∼1.75 times more) treated culture, while no significant difference was detected in the HL treated condition (Fig. 8C). The two-way ANOVA analysis indicates that for proteins both factors (growth treatment and light regime) and their interaction significantly influenced this parameter. Conversely, carbohydrates and lipids were significantly influenced only by the growth treatment (Supplementary Table 2 and 3).

**Figure 8.**
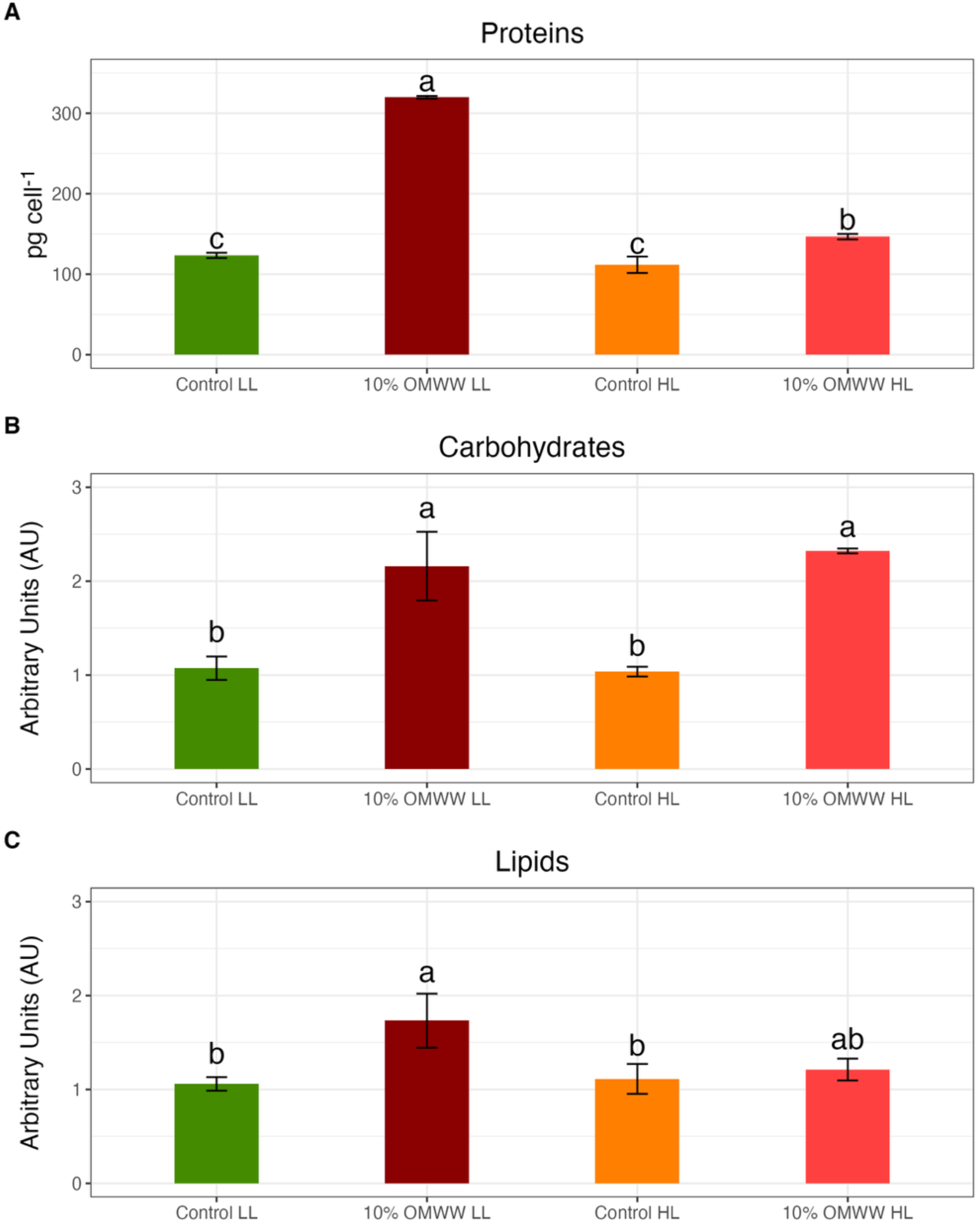
Macromolecular pools of *D. salina* at Low Light (LL) and High Light (HL) in 10% OMWW. Quantification of (**A**) proteins (pg cell^−1^), semiquantification of (**B**) carbohydrates (AU) and (**C**) lipids (AU) of *D. salina* in LL control cultures (green), 10% OMWW LL (bordeaux), HL control cultures (orange) and 10% OMWW HL (orange) cultures during the exponential growth phase. Different letters represent statistical differences between groups resulting from two-way ANOVA analysis (factors: growth condition and light regime).

#### 3.2.4 Reduction state of the biomass

The 10% OMWW treated culture showed different reduction indexes when exposed to different light conditions (Fig. 9). At LL, the biomass had a lower reduction index compared to the control, while at HL the reduction index of the biomass was higher than the control conditions. The two-way ANOVA analysis indicates that the light regime and its interaction with the growth condition significantly influenced this parameter (Supplementary Table 2 and 3).

**Figure 9.**
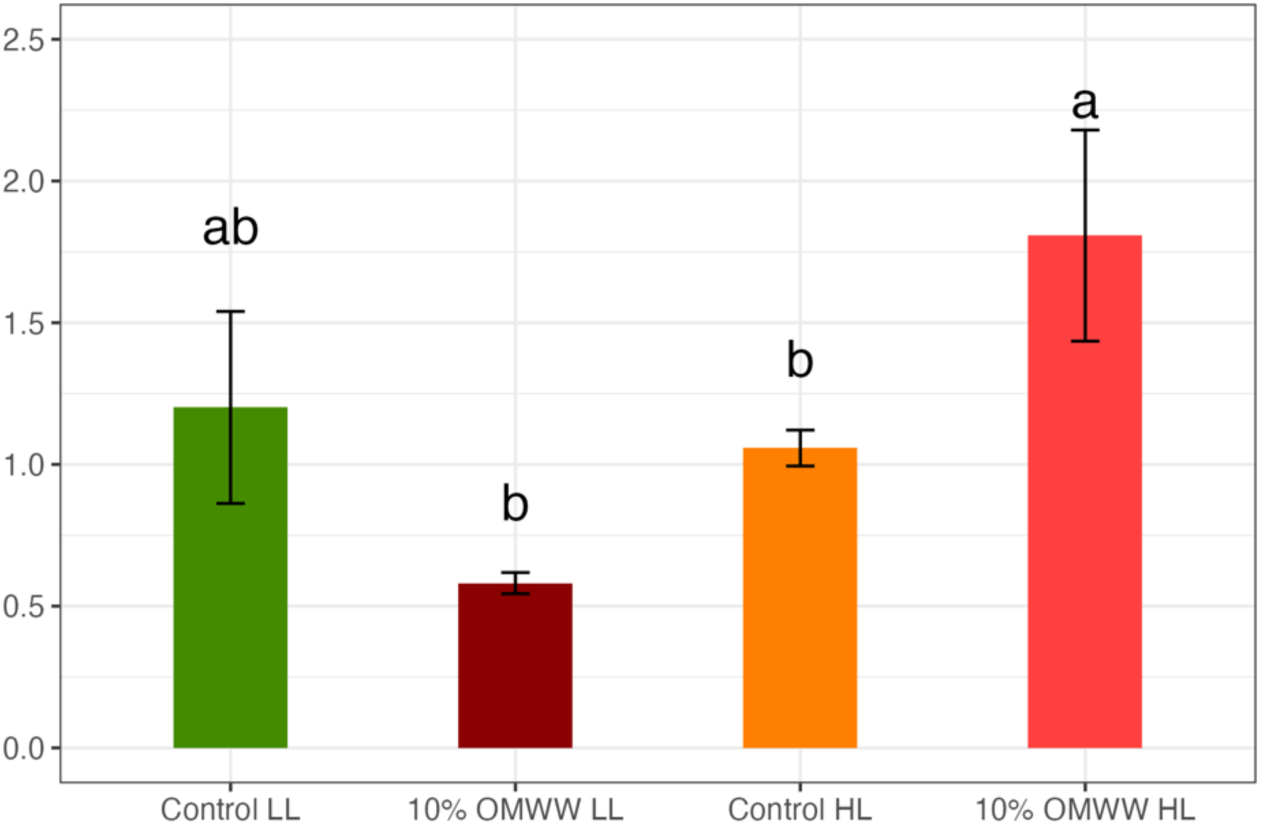
Reduction index of *D. salina* at Low Light (LL) and High Light (HL) in 10% OMWW. Quantification of the reduction index of *D. salina*’s biomass in LL control cultures (green), 10% OMWW LL treated cultures (bordeaux), HL control cultures (orange) and 10% OMWW HL treated cultures (orange) during the exponential growth phase. Different letters represent statistical differences between groups resulting from two-way ANOVA analysis (factors: growth treatments and light regime).

## 4. Discussions

### 4.1 Oxidative stress and genotoxic damages induced by phenolic compounds during short-term exposure

Exposure to 10% OMWW induced a marked increase in the DPPH radical-scavenging activity of *D. salina* extracts after 2 days, compared with the corresponding control (Fig. 2). Previous studies have similarly reported enhanced antioxidant capacity in microalgae cultivated in olive mill wastewater. In particular, Dahmen-Ben Moussa et al. (2021) observed increased DPPH scavenging activity in *Scenedesmus* sp. grown in OMWW, associated with the accumulation of phenolic compounds in the algal biomass. However, while their study primarily focused on the antioxidant potential of biomass obtained under different cultivation strategies, the present work investigated temporal changes in antioxidant capacity following exposure to OMWW and individual phenolic compounds.

Importantly, the DPPH assay measures the radical-scavenging capacity of extracts *in vitro* and does not directly quantify intracellular oxidative stress. Therefore, the increased DPPH inhibition observed after 2 days may reflect changes in cellular antioxidant defences in response to OMWW exposure, although a contribution from OMWW-derived phenolic compounds associated with the biomass cannot be excluded.

The early increase in DPPH scavenging activity was accompanied by genotoxic effects detected through the Comet assay after 24 h of exposure. In particular, OMWW-treated cells exhibited significantly higher Tail Intensity (TI) and Tail Moment (TM) values compared with several control and treatment conditions (Figs. 4 and 5). Although the two assays were performed at different time points, these findings suggest that OMWW exposure was associated with early alterations in antioxidant capacity and DNA integrity. Adverse effects of OMWW exposure have also been reported in other aquatic organisms, including *Mytilus galloprovincialis* and *Unio tumidiformis* (Danellakis et al., 2011; Costa et al., 2026), further highlighting the potential toxicity of these effluents.

Among the individual phenolic compounds tested, tyrosol showed a notable temporal variation in genotoxic parameters, with significant decreases in Tail Length (TL), TI and TM between the first and eighth days of exposure (Figs. 3–5). These changes suggest a stronger initial genotoxic response that progressively declined over time. Since tyrosol is one of the most abundant phenolic compounds in OMWW (Babić *et al*., 2019), it may contribute to the genotoxic effects observed following OMWW exposure. Nevertheless, the complex composition of OMWW prevents a direct attribution of its toxicity to tyrosol alone. Previous investigations on *D. salina* exposed to phenol reported alterations in nucleolar ultrastructure (Duan et al., 2017), while genotoxic effects of phenolic compounds have also been described in other biological models (Da Silva et al., 2002; Rocha-Guzmán et al., 2009; Azqueta and Collins, 2016).

Interestingly, antioxidant and genotoxic responses changed substantially with increasing exposure time. After 7 days, DPPH inhibition in OMWW-treated cultures significantly decreased compared with the initial measurement and was no longer significantly different from the corresponding control (Fig. 2). A significant reduction in DPPH inhibition over time was also observed in cultures exposed to caffeic acid and the phenolic mixture, whereas tyrosol-treated cultures showed an opposite trend. Similarly, the genotoxic response observed during early exposure progressively decreased. After 8 days, TM values no longer differed significantly among treatments, while TI in OMWW-treated cultures was no longer significantly different from its corresponding HL control, although differences relative to Control LL and Mix persisted (Figs. 3–5). These findings are consistent with complete recovery of the early OMWW-associated genotoxic response relative to the corresponding control, particularly in terms of TM, suggesting that *D. salina* can effectively cope with the initial effects of OMWW exposure. Moreover, growth rates were not significantly affected by the tested treatments, although cultures exposed to the phenolic mixture exhibited a lower final cell concentration than the LL control (Fig. 1; Table 1). Overall, these observations support a remarkable capacity of *D. salina* to tolerate phenolic compounds and OMWW, potentially through physiological acclimation mechanisms, in agreement with previous evidence of its tolerance to phenolic compounds (Mollo et al., 2023).

Although antioxidant and genotoxic responses to environmental stressors have been investigated in several microalgal species, studies integrating early responses and subsequent physiological changes over time remain relatively scarce. Evaluations of antioxidant capacity using the DPPH assay have frequently focused on final endpoints after prolonged exposure to pollutants (Danouche et al., 2020) or measurements following several days of treatment (Dahmen-Ben Moussa et al., 2021; Cao et al., 2023; Santos et al., 2025). Other studies have investigated temporal variations in antioxidant activity under different experimental contexts. For example, Aremu et al. (2014) monitored changes in DPPH inhibition after 5 and 10 days in five *Scenedesmus* strains using different extraction methodologies to assess the influence of culture age, while Ismail and El Zokm (2023) evaluated variations in DPPH radical-scavenging activity within a natural phytoplankton community subjected to heavy metal pollution. Conversely, microalgal studies employing the Comet assay have often focused on relatively short exposure periods, ranging from a few hours to 24 h (Aoyama et al., 2003; Prado et al., 2015; Chavan et al., 2024), although longer-term assessments of DNA damage have also been reported (Cheng et al., 2024). Even in *Dunaliella* species, genotoxic assessments have generally been restricted to narrow temporal windows (Barmshuri et al., 2023). This methodological fragmentation highlights the value of integrating antioxidant and genotoxic measurements at different stages of exposure to better characterize microalgal responses to environmental stressors.

Overall, exposure to 10% OMWW induced pronounced early changes in DPPH radical-scavenging activity and genotoxic parameters in non-acclimated *D. salina*. Notably, these responses were not fully reproduced by exposure to the mixture of selected phenolic compounds, suggesting that the biological effects of OMWW cannot be explained solely by the three phenolics investigated. Other constituents of the wastewater, as well as interactions among its components, may contribute to the observed responses. However, because OMWW and individual phenolic treatments were tested under different light regimes, their effects cannot be directly compared without considering the influence of irradiance. Nevertheless, the subsequent decrease in DPPH inhibition, together with the complete recovery of the early TM response by day 8, supports the ability of *D. salina* to overcome the initial disturbance induced by OMWW. These findings highlight the pronounced tolerance of this species and suggest that physiological acclimation contributes to its resilience under the experimental conditions tested.

### 4.2 *D. salina* growth in OMWW is favoured under a High Light regime

Growth of *D. salina* cells acclimated to 10% OMWW medium at LL was still significantly slower than in the control: division time was three times longer and maximum density about three times lower (Fig. 6). The activation of antioxidant defences in response to the pro-oxidant activity of phenolic compounds (Scalbert *et al.,* 2005; Abbas *et al.,* 2017), also reported for *Lingulodinium polyedrum* (Martins *et al.,* 2015), likely imposed an additional energetic burden on *D. salina*. This metabolic cost may explain the reduced growth performance observed under OMWW exposure. The partial recovery under high-light conditions suggests that increased photosynthetic energy availability compensated, at least in part, for the resources allocated to stress defence and cellular maintenance.

Despite this improved growth, concentration of phenolic compounds in the medium did not change during the experiment (Supplementary Fig. 6). Microalgae can remove phenolics from wastewater by mineralization, biotransformation and biosorption (Lindner and Pleissner, 2019; Mollo *et al.,* 2026). However, the lack of a cell wall limits biosorption in *D. salina* (Mollo *et al.,* 2026), and the absence of any change in phenolic content indicates that no other removal mechanism operates, in line with the results for single phenolics in this species (Mollo *et al.,* 2023). Genetic engineering, already applied in *Dunaliella* (Hassanien *et al.,* 2023; Ambily *et al.,* 2025), or Adaptive Laboratory Evolution (ALE), which improved phenol removal in the marine microalga *Isochrysis galbana* (Li *et al.,* 2021), could overcome this limitation. Since *I. galbana* showed no adsorption capacity (Mollo *et al.,* 2026), ALE may induce different removal mechanisms, like an increase of phenol hydroxylase activity, as observed for *I. galbana* (Li *et al.,* 2021), or mineralisation and/or bioaccumulation.

The response of the photosynthetic apparatus further supports the hypothesis that OMWW imposed a substantial physiological burden on *D. salina*. Under LL condition, the reductions in chlorophylls, carotenoids and the Chl *a*/Chl *b* ratio are consistent with oxidative stress-mediated alterations of the photosynthetic machinery, as ROS are known to damage photosystems and pigment-protein complexes (Sabatini *et al.,* 2009; Zhang *et al.,* 2015). In contrast, although chlorophyll content remained lower in OMWW-grown cells under HL, pigment composition was maintained closer to control values, indicating a more balanced organization of the photosynthetic apparatus.

The improved physiological performance observed at high irradiance suggests that the negative effects of OMWW were primarily related to the metabolic costs associated with stress mitigation rather than to limitations in light availability. Indeed, previous studies have shown that the absorption spectrum of OMWW does not substantially overlap with the major absorption peaks of chlorophylls *a* and *b* (Elhajjouji *et al.,* 2007; Rizzo *et al.,* 2008). Therefore, the reduced growth observed under low light is more likely attributable to insufficient energy availability to simultaneously sustain cellular maintenance and stress-response mechanisms. By increasing irradiance, *D. salina* was able to allocate sufficient photosynthetic energy to both acclimation and growth, resulting in growth performances comparable to those of the control cultures. From an applied perspective, cultures grown in 10% OMWW under high light also displayed a favourable biochemical composition, characterized by elevated protein and carbohydrate contents, highlighting the potential of this cultivation strategy for biomass production and food-related applications (Fig. 8).Interestingly, we also observed a shift in the reduction index of the biomass (Fig. 9), which was higher in *D. salina* biomass at HL, suggesting a higher energy content (important for possible nutritional uses) in this growth condition (Fazeli Danesh *et al.,* 2017), compared to LL. The general increase of proteins, carbohydrates and lipids might be explained by a mixotrophic growth (Park *et al.,* 2012) of *D. salina* in 10% OMWW, as already observed in other chlorophytes such as *Chlorella sorokiniana* and other *Chlorella* species (Nirmalakhandan *et al.,* 2019; Hachicha *et al.,* 2025; Menaa *et al.,* 2025). Indeed, the COD value (chemical oxygen demand) of OMWW is generally high (Danellakis *et al.,* 2011; Sforza *et al.,* 2018). However, it is unclear whether *D. salina* can use organic carbon from the medium. Some studies hint at it (Capa-Robles *et al.,* 2021; Gonabadi *et al.,* 2022), but Celente *et al*. (2022) showed that the congeneric *D. tertiolecta* cannot, and depends on symbiotic bacteria for mixotrophy. Future experiments should test whether *D. salina* is mixotrophic in OMWW-based medium.

## 5. Conclusions

In conclusion, *D. salina* showed a remarkable capacity to acclimate to the oxidative and genotoxic stress induced by OMWW, particularly under high-light conditions, where growth performance was substantially improved. Although *D. salina* appears suitable for cultivation in OMWW, its limited capacity to remove phenolic compounds suggests that this species may be more valuable for biomass production than as a stand-alone agent for OMWW phycoremediation.

## Acknowledgments

None to declare.

## Conflict of interest

The authors declare that the research was conducted in the absence of any commercial or financial relationships that could be construed as a potential conflict of interest.

## CRediT authorship contribution statement

**Filippo Drigo**: Conceptualization, Formal analysis, Investigation, Visualization, Writing – original draft, Writing – review & editing. **Lorenzo Mollo**: Conceptualization, Methodology, Visualization, Supervision, Writing – original draft, Writing – review & editing. **Fabio Marcheggiani**: Methodology, Formal analysis, Writing – review & editing. **Luca Tiano**: Methodology, Supervision, Writing – review & editing. **Alessandra Norici**: Conceptualization, Methodology, Supervision, Writing – review & editing.

## Funding

There are no fundings to be declared.

## Data Availability

Data will be available upon request.

## Notes

### Competing Interest Statement

The authors have declared no competing interest.

