## Supplementary Figures and Supplementary Table 2 for "Acclimation of *Dunaliella salina* to Olive Mill Wastewater: Oxidative Stress, Genotoxicity and the Role of Light Availability"

**Supplementary material**

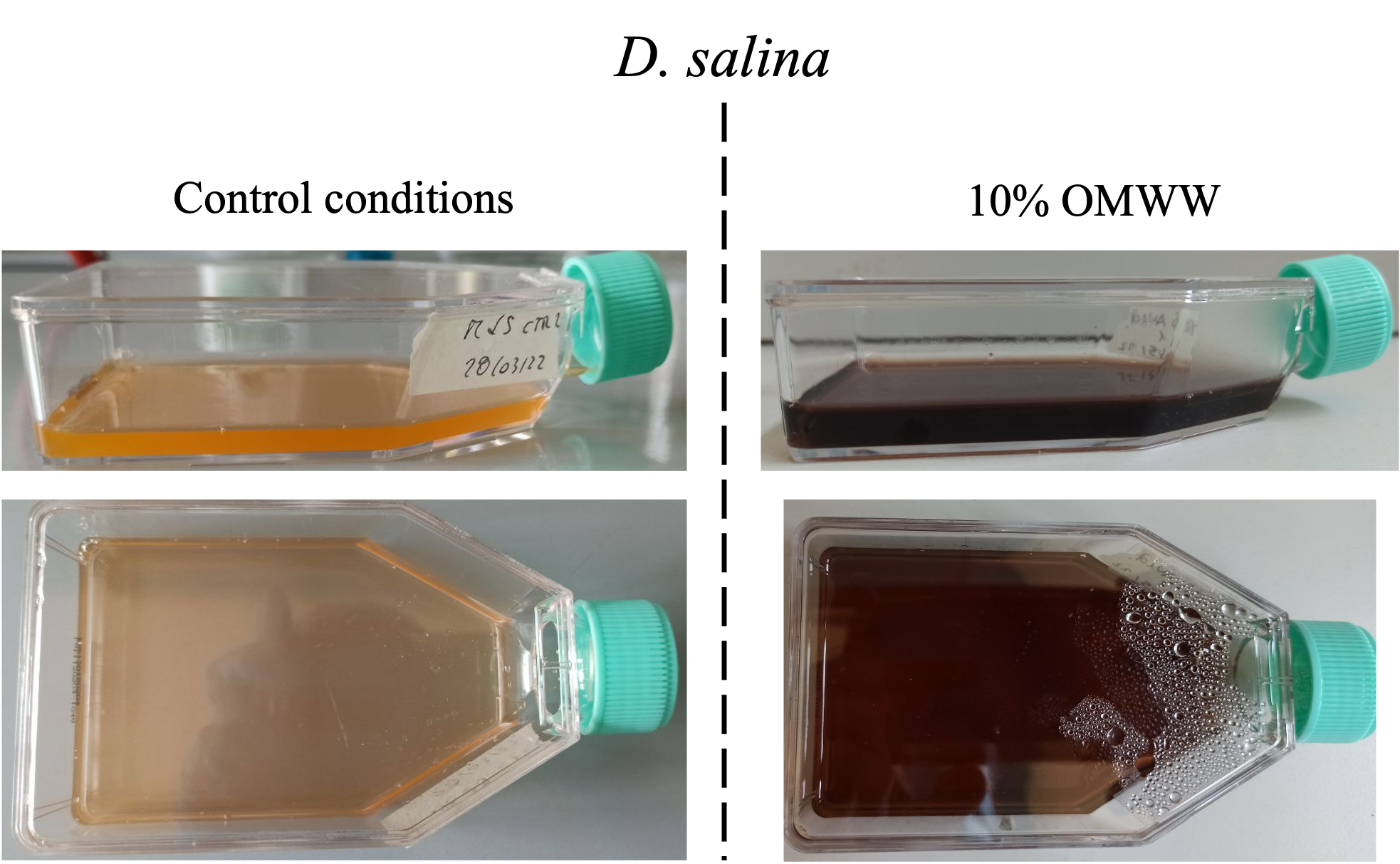

**Supplementary Figure 1. Representation of *D. salina* in control conditions and in 10% OMWW.** Top and side images showing the difference in color of *D. salina* cultures in control conditions at the end of the exponential growth phase and in 10% OMWW, highlighiting the dark color of this medium.

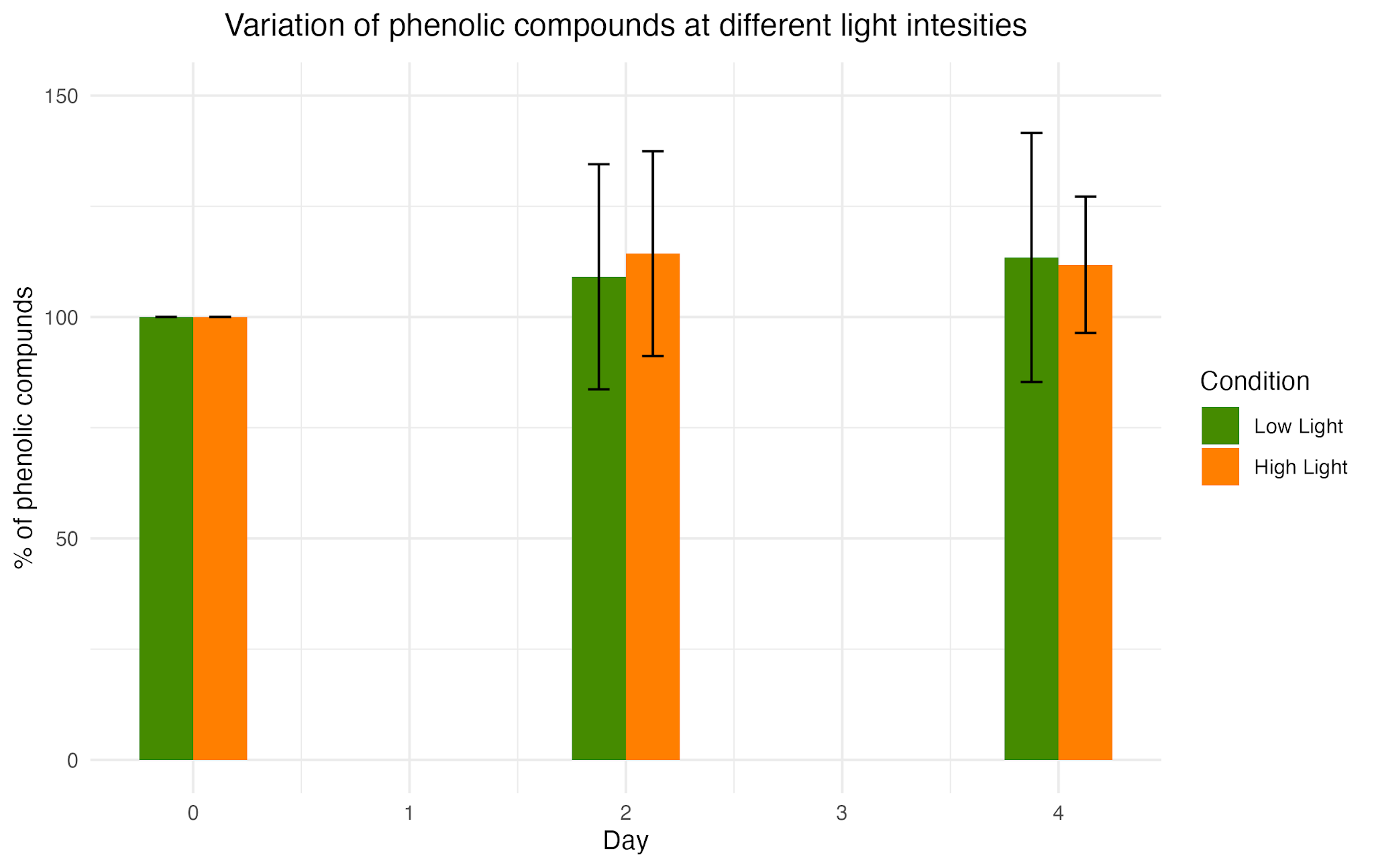

**Supplementary Figure 2. Variation of phenolic compounds during growth of *D. salina*.** Variation of phenolic compounds expressed as percentage of initial concentration in the OMWW at LL (green) and HL (orange) until the middle of the exponential phase (day 4).

**Supplementary Table 1** Summary of one-way ANOVA Tukey's post hoc test. Results of Tukey's post hoc test for significant findings conducted on *D. salina* for the genotoxic effects analysis of OMWW**.**

***Available as a separate .xlsx file***

**Supplementary Table 2.** Summary of two-way ANOVA analysis. Results of two-way ANOVA analysis for factors effects and interaction conducted on *D. salina* for the OMWW growth characterization.

| **Two-way ANOVA** | | |  | **Df** |  | **Sum.Sq** |  | **Mean.Sq** |  | **F.value** |  | **Pr..F.** |  | **Figure** |
| --- | --- | --- | --- | --- | --- | --- | --- | --- | --- | --- | --- | --- | --- | --- |
| **Parameter** |  | **Factors** |  |  |  |  |  |  |  |  |  |  |  |  |
| Chlorophyll *a* |  | growth_condition |  | 1 |  | 7.81983075 |  | 7.81983075 |  | 95.1900484 |  | 1.02E-05 |  | 7A |
|  |  | light_regime |  | 1 |  | 1.61700208 |  | 1.61700208 |  | 19.683611 |  | 0.00217772 |  |  |
|  |  | growth_condition:light_regime |  | 1 |  | 0.16921875 |  | 0.16921875 |  | 2.05988359 |  | 0.18914005 |  |  |
|  |  | residuals |  | 8 |  | 1 |  | 0.08214967 |  | NA |  | NA |  |  |
| Chlorophyll *b* |  | growth_condition |  | 1 |  | 0.47880075 |  | 0.47880075 |  | 58.2234754 |  | 6.13E-05 |  | 7B |
|  |  | light_regime |  | 1 |  | 0.30369008 |  | 0.30369008 |  | 36.9295414 |  | 0.00029692 |  |  |
|  |  | growth_condition:light_regime |  | 1 |  | 0.00110208 |  | 0.00110208 |  | 0.13401634 |  | 0.72379535 |  |  |
|  |  | residuals |  | 8 |  | 0.065788 |  | 0.0082235 |  | NA |  | NA |  |  |
| Chlorophyll *a*/  Chloprophyll *b* |  | growth_condition |  | 1 |  | 0.18700033 |  | 0.18700033 |  | 8.10323298 |  | 2.16E-02 |  | 7C |
|  |  | light_regime |  | 1 |  | 0.266412 |  | 0.266412 |  | 11.5443565 |  | 0.00939238 |  |  |
|  |  | growth_condition:light_regime |  | 1 |  | 0.08840833 |  | 0.08840833 |  | 3.83097351 |  | 0.086018 |  |  |
|  |  | residuals |  | 8 |  | 0.184618 |  | 0.02307725 |  | NA |  | NA |  |  |
| Carotenoids |  | growth_condition |  | 1 |  | 29.2875008 |  | 29.2875008 |  | 14.8069932 |  | 4.89E-03 |  | 7D |
|  |  | light_regime |  | 1 |  | 0.26137008 |  | 0.26137008 |  | 0.13214187 |  | 0.72564018 |  |  |
|  |  | growth_condition:light_regime |  | 1 |  | 1.41796875 |  | 1.41796875 |  | 0.71688785 |  | 0.42177059 |  |  |
|  |  | residuals |  | 8 |  | 15.8236047 |  | 1.97795058 |  | NA |  | NA |  |  |
| Proteins |  | growth_condition |  | 1 |  | 29107.8994 |  | 29107.8994 |  | 1047.18521 |  | 5.29E-07 |  | 8A |
|  |  | light_regime |  | 1 |  | 15935.3458 |  | 15935.3458 |  | 573.289682 |  | 2.3675E-06 |  |  |
|  |  | growth_condition:light_regime |  | 1 |  | 14245.4231 |  | 14245.4231 |  | 512.493057 |  | 3.1264E-06 |  |  |
|  |  | residuals |  | 8 |  | 138.98162 |  | 27.796324 |  | NA |  | NA |  |  |
| Carbohydrates |  | growth_condition |  | 1 |  | 3.40078423 |  | 3.40078423 |  | 67.7918583 |  | 1.73E-04 |  | 8B |
|  |  | light_regime |  | 1 |  | 0.00936407 |  | 0.00936407 |  | 0.18666512 |  | 0.68080179 |  |  |
|  |  | growth_condition:light_regime |  | 1 |  | 0.02347097 |  | 0.02347097 |  | 0.46787473 |  | 0.51949643 |  |  |
|  |  | residuals |  | 8 |  | 0.3009905 |  | 0.05016508 |  | NA |  | NA |  |  |
| Lipids |  | growth_condition |  | 1 |  | 0.49172826 |  | 0.49172826 |  | 13.7109708 |  | 1.01E-02 |  | 8C |
|  |  | light_regime |  | 1 |  | 0.13090709 |  | 0.13090709 |  | 3.65011218 |  | 0.10463148 |  |  |
|  |  | growth_condition:light_regime |  | 1 |  | 0.19734278 |  | 0.19734278 |  | 5.50255366 |  | 0.05739073 |  |  |
|  |  | residuals |  | 8 |  | 0.21518312 |  | 0.03586385 |  | NA |  | NA |  |  |
| Reduction index |  | growth_condition |  | 1 |  | 0.012616 |  | 0.012616 |  | 0.19494291 |  | 6.71E-01 |  | 9 |
|  |  | light_regime |  | 1 |  | 0.87927955 |  | 0.87927955 |  | 13.5866573 |  | 0.00616554 |  |  |
|  |  | growth_condition:light_regime |  | 1 |  | 1.40664218 |  | 1.40664218 |  | 21.7354825 |  | 0.00161896 |  |  |
|  |  | residuals |  | 8 |  | 0.5177312 |  | 0.0647164 |  | NA |  | NA |  |  |

**Supplementary Table 3.** Summary of two-way ANOVA Tukey's post hoc. Results of Tukey's post hoc test for significant findings conducted on *D. salina* for the OMWW growth characterization.

***Available as a separate .xlsx file***
